# Slow phase-locked endogenous modulations support selective attention to sound

**DOI:** 10.1101/2021.02.03.429516

**Authors:** Magdalena Kachlicka, Aeron Laffere, Fred Dick, Adam Tierney

## Abstract

To make sense of complex soundscapes, listeners must select and attend to task-relevant streams while ignoring uninformative sounds. One possible neural mechanism underlying this process is alignment of endogenous oscillations with the temporal structure of the target sound stream. Such a mechanism has been suggested to mediate attentional modulation of neural phase-locking to the rhythms of attended sounds. However, such modulations are compatible with an alternate framework, where attention acts as a filter that enhances exogenously-driven neural auditory responses. Here we attempted to adjudicate between theoretical accounts by playing two tone steams varying across condition in tone duration and presentation rate; participants attended to one stream or listened passively. Attentional modulation of the evoked waveform was roughly sinusoidal and scaled with rate, while the passive response did not. This suggests that auditory attentional selection is carried out via phase-locking of slow endogenous neural rhythms.

## Introduction

As we navigate through the world, we are bombarded by more sensory information than we can process and respond to. One of the brain’s most important tasks, therefore, is to focus on the most relevant information while ignoring the rest. Whereas the eye can be directed to point at and focus on regions of visual space, this mechanical strategy is generally not available to the human auditory system. Instead, it must use neural mechanisms to segregate acoustic information into sources or ‘streams’, select one particular stream, and then actively attend to that stream to extract task-relevant information (Shinn-Cunningham, 2008; Holt et al., 2018).

What neural strategies might the listener use to focus in on and track a task-relevant stream or signal? One possibility is to winnow out or filter auditory streams based on acoustic dimensions along which they differ. For example, if one stream lies in a higher frequency band than the other, listeners could selectively attend to that stream by directing attention to a particular frequency band (spectrally-selective attention; Fritz et al., 2012; Da Costa et al., 2013; Dick et al. 2017; Riecke et al., 2017). Another possible strategy is to take advantage of timing or rhythmic differences between streams, so that attention can be directed to time points likely to contain the to-be-attended stream and less likely to contain the to-be-ignored stream (Nobre & van Ede, 2018).

That listeners make use of temporally-selective attention to select sound streams for further processing is supported by findings that temporal differences between a target stream and a distractor stream help boost performance on attention tasks. Prior knowledge about target timing onsets has been shown to boost tone detection (Bonino & Leibold, 2008), birdsong identification (Best et al., 2007), word recognition (Gatehouse & Akeroyd, 2008), and speech comprehension (Kitterick et al., 2010) when target stimuli are presented among distractors. Indeed, when speaking, talkers modulate the temporal characteristics of speech to minimize overlap with background speech, thus allowing listeners to use such a temporally-selective strategy to understand what the speaker is saying (Cooke & Lu, 2010).

How might the brain carry out such temporally-selective attention? One possibility is that brain activity “entrains” to the attended stream, such that endogenous neural oscillations phase-lock to its temporal structure (Schroeder & Lakatos, 2009). Indeed, when non-human primates attend to one stream in an inter-modal attention task, slow neural activity aligns with the temporal structure of the attended stream (Lakatos et al., 2008). Similarly, when humans undergoing intracranial recording for epilepsy performed an inter-modal attention task with alternating auditory and visual stimuli (Besle et al., 2011), attention to one versus the other modality was linked to a shift in phase of 180 degrees in a slow modulation of neural activity at the within-modality presentation rate.

This neural entrainment hypothesis has influenced a large body of cognitive neuroscience research on selective attention. However, the vast majority of this research does not directly examine the exact form of the attentional modulation of neural activity. Instead, most studies rely on less direct measures of neural entrainment, such as inter-trial phase-locking. For example, Lakatos et al. (2013) showed that when rhesus macaques attend to one of two tone streams presented at different rates, this leads to increased phase-locking at the attended rate. A further study with macaques (Lakatos et al., 2016) demonstrated that periods with a high degree of phase-locking at the attended stimulus rate were associated with greater hit rates, while decreased phase-locking occurred during periods with greater false alarm rates along with increased alpha-band amplitude (suggesting an increase in inattention – Madore et al., 2020).

One of the advantages of using intertrial phase-locking and phase alignment to study attentional modulation of neural entrainment is that data can be acquired non-invasively in human auditory EEG or MEG auditory attention paradigms. For example, human participants’ phase-locking to a task-relevant sound stream presented in an environmental noise background has been shown to increase after the presentation of a target and decrease after salient distractors (Huang & Elhilali, 2020). When participants attend to one of two temporally interdigitated tone streams there is a roughly 180-degree shift in the phase of the neural response at the tone stream presentation rate (Laffere et al., 2020a, b). Studies of selective attention to speech have shown that low-frequency neural activity more closely mirrors the slow amplitude modulation patterns of the stimulus stream being attended, compared to the stream being ignored (Kerlin et al., 2010; Ding & Simon, 2013; Zion Golumbic et al., 2013; O’Sullivan et al., 2015; Ghinst et al., 2016). This phenomenon is sometimes interpreted according to the neural entrainment framework, i.e. as reflecting entrainment of cortical oscillations to the temporal structure of attended speech (Horton et al., 2013; Riecke et al., 2018; Fuglsang et al., 2020; for a discussion, see Obleser & Kayser, 2019).

Such research has shown that attention modulates the strength of the relationship between neural activity and the temporal structure of the attended signal. However, this entrainment in a broad sense—the presence of stimulus-brain temporal alignment—is not strong evidence for entrainment in a narrow sense. As defined by Obleser and Kayser 2019, a more stringent test of entrainment in a narrow sense is to show that the phase of endogenous, ongoing oscillators is adjusted, such that peaks align with timepoints containing acoustic edges in attended stimuli. With the exception of Lakatos et al. 2008 and Besle et al. 2011, both of whom showed evidence for the existence of slow, sinusoidal neural rhythms aligned with the temporal structure of attended stimulus streams, the remainder of these findings are consistent with an alternate explanation – namely that attention acts as a filter attenuating and amplifying exogenous responses to sound.

Attentional modulation of the magnitude of responses (ERPs to sound onset or amplitude-envelope-following responses) could potentially be observed as increases in phase-locking, or shifts in neural phase at the rate of stimulus presentation, thus leading to the results summarized above. Indeed, a large body of research has shown that selective attention to a sound stream can increase the amplitude of exogenously-evoked potentials to sound onsets within the target stream (Hillyard et al., 1973; Chait et al., 2010; Choi et al., 2013; Dai et al., 2018), including the P50 and N100 components (Woldorff et al., 1993). Moreover, prior knowledge about the likely time of onset of a stimulus can also enhance the N100 (Lange et al., 2003), suggesting that attention to time could modulate the amplitude of exogenous responses. Thus, temporal structure in an attended stimulus could set up a reliable expectation of future stimulus onsets. This in turn would enhance responses to these stimuli, and lead to the increased phase-locking commonly reported as a neural correlate of auditory selective attention.

In sum, much of the existing research on auditory selective attention is consistent with two competing explanations: the first positing selective attention as a filter that amplifies or attenuates exogenous responses to stimuli, and the other suggesting that endogenous oscillators synchronize to the temporal structure of attended stimuli. In the study reported below, we designed an EEG experiment to potentially adjudicate between these two explanations. Two groups of participants were asked to attend to one of two streams composed of temporally interleaved tone sequences. Across conditions, we manipulated tone presentation rate; across groups, we varied tone duration. By examining the shape of the attentional modulation of the ERP waveform, we can test several predictions linked to the neural entrainment versus attentional filter accounts. First, if the ‘attention as neural filter’ account holds, and attention modulates the amplitude of event-related responses to sound onset, then the width of the attentional modulation should be roughly constant across presentation rate conditions. In particular, the attentional modulation should be limited to the time window of the N100 – or, if it extends to time points containing adjacent positive components, the modulation’s polarity should flip (changing from negative to positive). Alternately, if attention modulates the amplitude of envelope-following responses, the width of the modulation should scale with tone duration. On the other hand, the neural entrainment account suggests that the attentional modulation should scale more or less linearly with rate, such that modulated portions of the waveform become longer at slower rates, but be relatively unaffected by tone duration. Finally, the neural entrainment account predicts that attentional modulations should continue through the silence between tone sequences, as oscillations are hypothesized to be somewhat self-sustaining (Large & Kolen, 1994). By contrast, the attentional filter account predicts that attentional modulations should die out rapidly after the offset of the final tone of the sequence.

## Materials and Methods

### Participants

Participants were postgraduate students enrolled in programmes related to auditory processing (audiology, auditory neuroscience and acoustics) or professional musicians or audio engineers living in London. Participants were selected for higher levels of auditory experience and expertise because the experiment was quite demanding of auditory attention, and in previous studies, non-expert participants required somewhat more training to achieve good performance in the task (Laffere et al., 2020a). 15 participants (aged 19-48, M=29.47, SD=7.41; 9 females) took part in the first experiment (with variable tone durations across presentation rates). 14 different participants (aged 22-36, M=28.36, SD=4.89; 9 females) took part in the second experiment (with a single short fixed tone duration across presentation rates). All participants reported no prior diagnosis of hearing impairment or neurological disorders affecting hearing. The experimental paradigm was approved by the Research Ethics Committee of the Department of Psychological Sciences at Birkbeck, University of London.

### Experimental and stimulus design and presentation

In both experiments, participants listened to series of interleaved tone sequences segregated into high and low frequency bands (see Figure 1). Both experiments used a fully within-subject design, crossing attention condition (attend high band sequences, attend low band sequences, listen passively) with within-band tone presentation rate (2Hz, 3Hz, 4Hz, 5Hz). Each of the 12 combinations of attention condition by tone presentation rate was completed by each participant, with condition order randomized across participants.

**Figure 1.**
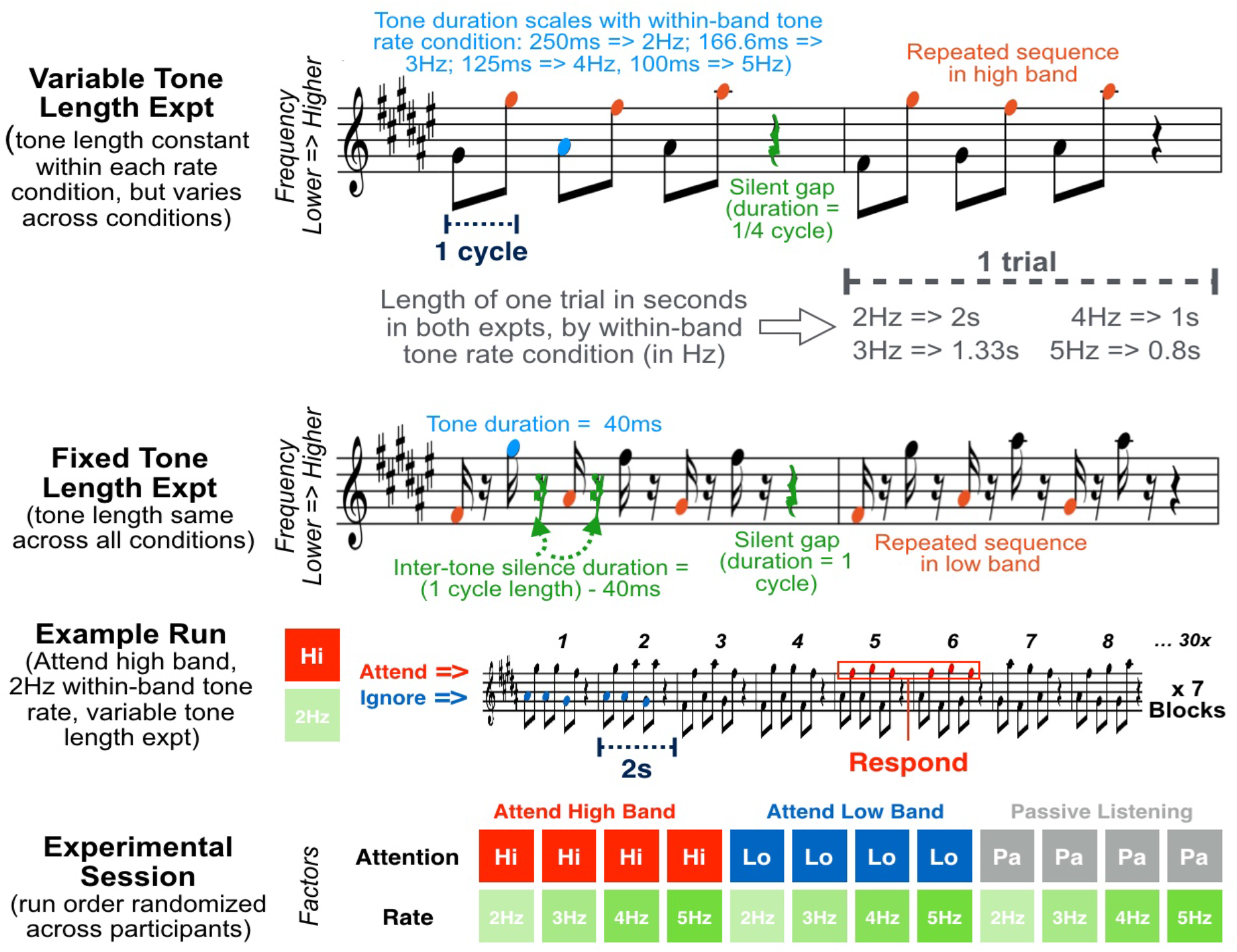
Schematics showing experimental design. The top row shows two trials of the Variable Tone Length Experiment, and the second row two trials of the Fixed Tone Length Experiment, in quasi-musical notation. For ease of depiction on one stave, fundamental frequencies depicted by musical notes are one octave higher than those used in the experiment. For each tone rate condition in the variable tone length experiment, all tones are set to a condition-specific duration (see Figure and Methods); in the fixed tone length experiment, all tones are 40ms long regardless of rate condition. The third row shows a segment of an example run of the ‘attend high band, 2Hz within-band tone presentation rate’ condition from the variable tone length experiment. Each run is made up of 7 blocks of 30 trials, where a trial contains two interleaved 3-tone sequences, one in the higher frequency band, and one in the lower band. There are three high/low-tone ‘cycles’ in a trial, followed by a silence equal in duration to an additional cycle. The bottom row shows all twelve runs for a single experiment, where each run is a combination of attention condition and within-band tone presentation rate. Note that run order is randomized across participants. See attached files on biorxiv for example blocks.

The basic stimulus units were cosine-ramped pure tones constructed at a 48 kHz sampling rate using MATLAB (The MathWorks, Inc., Natick, MA). These tones were arranged into sequences of six, followed by a silent period; tone and silence duration varied with experiment and condition, as explained below. Stimuli were presented diotically through Etymotic 3A insert earphones (Etymotic, Elk Grove Village, IL) at an 80-dB sound pressure level.

The tones were presented in two frequency bands; for each band, a set of three possible fundamental frequencies (185, 207.7, and 233.1 Hz for the low band and 370, 415.3, and 466.2 Hz for the high band) was randomly sampled to create mini-sequences of three tones. These tones were presented 180 degrees out of phase with a within-band rate of presentation varying between blocks of 2 Hz, 3 Hz, 4 Hz, or 5 Hz (i.e. with 500 ms, 333 ms, 250 ms or 200 ms between tone onsets respectively). Tones in low (A) and high (B) frequency bands alternated, forming a repeating ABABAB pattern followed by a silence equal in duration to one cycle of the within-band rate (i.e. 500 ms, 333 ms, etc). The ABABAB pattern plus silence composed one trial. (Note that the first tone of each sequence was always a low tone; this ensured that time of onset of sequences in each frequency band was always predictable, facilitating stream segregation and selection). Each block consisted of 30 trials; 3-6 sequence repetitions were spaced semi-randomly within each frequency band, with the constraint that there was at least one non-repeating sequence between the repetitions. Finally, for each of the two experiments, there was a single run for each condition, with seven 30-trial blocks per run.

The Variable Tone Length and Fixed Tone Length Experiments differed in the length of the tones used to construct the sequences. In the Variable Tone Length Experiment, tone length differed across the presentation rate conditions, such that, within each condition, tone duration was always equal to half of one cycle of the within-band presentation rate. This equalled 250 ms for 2 Hz, 166.66 ms for 3 Hz, 125 ms for 4 Hz, and 100 ms for 5 Hz. In the Fixed Tone Length Experiment, tones were always 40 ms in length, and this duration did not differ across the presentation rate conditions.

### Behavioural task

In the two ‘Active’ conditions, participants were asked to attend to tone sequences either in the low-or high-frequency band, while ignoring the tones presented in the competing frequency band. The participants’ task was to identify the repetition of a mini-sequence in the attended frequency band and respond by clicking the mouse. In a third “Passive” condition, participants were asked to sit quietly and listen to the tones without attending to either band, pressing a button at the end of each block to advance to the next block.

For the Active conditions, the latency window for recording behavioral responses was from (1/rate) s before to (5/rate) s after the end of the last tone in a given sequence. For example, for the 4 Hz condition, the window extended from 0.25 s before to 1.25 s after the end of the last tone in the sequence. Text feedback was displayed for correct answers, missed targets, and false alarms. Only one incorrect response could be recorded per sequence; in other words, two incorrect responses within the same sequence would be recorded as only a single false alarm. Performance was measured as d-prime, with the false alarm rate calculated using the total number of non-target sequences as the highest possible number of false alarms. A repeated measures ANOVA with one within-subjects factor (rate, levels 2, 3, 4, and 5 Hz) was used to investigate whether d-prime differed across rates.

### EEG data acquisition

Electrophysiological data were recorded using a BioSemi ActiveTwo 32-channel EEG system at a 16,384 Hz sample rate and with open filters in Acti-View (BioSemi) acquisition software. A standard 10/20 montage of active electrodes positioned in a fitted head cap with a sintered Ag-AgCl pallet was used. Two additional electrodes were placed on the left and right earlobes as external reference points. Contact impedance was kept below 20 kΩ throughout the testing session.

### EEG data processing

Event markers for the beginning of each block were recorded from trigger pulses sent to the neural data collection computer. The resulting data were downsampled to 500 Hz and eye blinks and muscle contraction artefacts were identified by independent component analysis and removed after visual inspection of component topographies and time courses. Data were segmented into epochs aligned with trial onsets, with epoch duration equaling 2.00, 1.33, 1.00, and 0.80 seconds for the 2, 3, 4, and 5 Hz conditions, respectively. Then, individual segments were excluded if the signal intensity of the channel exceeded ± 100 μV. All preprocessing steps were conducted with the use of custom MATLAB scripts and the FieldTrip M/EEG analysis toolbox (Oostenveld et al., 2011). Because we did not have an a priori hypothesis regarding the scalp location of effects, data were collapsed across all 32 electrodes prior to analysis.

Time-frequency analysis was conducted via a Hann-windowed FFT calculated over the entirety of each epoch. Inter-trial phase coherence (ITPC) was calculated at the frequency of within-band tone presentation by transforming the complex vector at this frequency to the unit vector (with a length of 1), averaging vectors across all trials, and then computing the length of the resulting vector as a measure of phase consistency. The resulting measure varies from 0 (no consistency in phase across trials) to 1 (perfect consistency in phase across trials). ITPC in the attend high and attend low conditions was averaged together and compared to ITPC in the passive condition to test the hypothesis that attention to one of the two bands would be linked to an increase in phase coherence at the attended rate, and that this attentional modulation would vary across frequencies. A 2 x 4 repeated measures ANOVA was run with two within-subjects factors, task (attention versus passive) and rate (2, 3, 4, and 5 Hz), with log-transformed ITPC at the frequency of tone presentation as the dependent variable.

The phase of the average vector was also extracted to examine the effects of attention on average phase. Given that high and low bands were presented 180 degrees out of phase, attention to one band versus the other should be linked to a phase difference across conditions (Laffere et al., 2020a,b). The effect of rate on the effect of attention on neural phase was investigated using a repeated measures ANOVA with one within-subjects factor (rate, 4 levels) and the dependent variable of the phase difference between attend high and attend low conditions.

Effects of attention on the event-related waveform were investigated by computing the average waveform across trials for the attend high, attend low, and passive conditions (separately for each presentation rate). Due to the continuous rhythmic stimulus presentation, there was not a meaningful pre-stimulus period for baseline amplitude calculation. Although there was a brief pause between sequences, one of our hypotheses was that effects of attention would continue through the pause. As a result, we could not assume that the period just before the onset of each trial was devoid of neural responses, and so epochs were baselined prior to averaging by subtracting the mean across the entire epoch.

Each sequence within each frequency band consisted of three tones. According to the neural entrainment hypothesis, therefore, each epoch should contain four cycles of a sinusoidal modulation of the ERP waveform due to attention: three aligned with the three tones of the sequence, followed by a fourth continuing through the silence. To test this hypothesis, we calculated the average passive waveform and attentional modulation waveform (attend-high minus attend-low waveforms) underlying a single cycle of the hypothesized sinusoid. For example, for the 5 Hz condition, we averaged together the period between 0 and 0.2, 0.2 and 0.4, and 0.4 to 0.6 seconds to calculate a single average response with a duration of 0.2 s. This period consisted of the average response to a pair of tones, a low tone (since the sequence in the low band always began first) followed by a high tone. To investigate the extent to which passive responses and attentional modulation continued into the silence between sequences, we separately analyzed the remaining portion of the response (which, in the 5 Hz condition, would be the period between 0.6 and 0.8 seconds).

For each of these waveforms – the average response to each ‘tone pair’ and the between-sequence silence – two separate sets of paired t-tests were conducted to determine the time points (2 ms temporal resolution) where 1) there was a significant difference in amplitude between the attend high and attend low conditions, and 2) the waveform in the passive condition was significantly different from zero. In addition, unpaired t-tests were conducted to determine the time points at which there was a difference between the short-tone and long-tone experiments in the attentional modulation (attend high minus attend low) and passive response. Each of these analyses was separately corrected for multiple comparisons using the False Discovery Rate method (Benjamini & Hochberg, 1995).

## Results

### Behavioral results

Attention performance (measured via d-prime and collapsed across Experiments 1 and 2, see **Table 1**) differed across the four presentation rates (F(3,84) = 19.81, p < 0.001), with better performance for slower rates. Post-hoc Bonferroni-corrected t-tests were used to examine pairwise differences in attention performance between rates. Performance was less accurate in the 4 Hz condition than the 2 Hz condition (t(28) = 3.86, p = 0.004). Performance was also less accurate in the 5 Hz condition compared to the 2 Hz (t(28) = 5.95, p < 0.001), 3 Hz (t(28) = 5.36, p < 0.001), and 4 Hz conditions (t(28) = 4.25, p = 0.001).

**Table 1.** Performance and neural metrics from time-frequency analyses across presentation rate conditions.

|  | 2Hz | 3 Hz | 4 Hz | 5 Hz |
| --- | --- | --- | --- | --- |
| **Attention performance (dprime)** | 2.83(0.79) | 2.74(0.81) | 2.50(0.81) | 2.17(0.88) |
| **Active log(ITPC)** | -1.92 (0.43) | -1.89 (0.38) | -2.02 (0.36) | -2.17 (0.27) |
| **Passive log(ITPC)** | -2.37 (0.31) | -2.46 (0.35) | -2.36 (0.43) | -2.37 (0.29) |
| **Attend high – low phase diff (radians)** | 1.87(0.98) | 2.32(0.78) | 2.11(0.89) | 1.51(0.98) |

### EEG time-frequency analyses

Because we did not have a prior hypothesis regarding effects of tone duration on attentional modulation of either ITPC or neural phase, EEG time-frequency analyses were collapsed over the Variable Tone Duration and Fixed Tone Duration experiments. When participants were asked to attend to one of the two bands and detect occasional repeated sequences, ITPC was higher at the within-band presentation rate than for the passive condition (main effect of task F(1,28) = 42.24, p < 0.001; see **Table 1**). Although there was no overall ITPC difference across rates (F(3,84) = 1.62, p > 0.1), the ITPC difference between active and passive conditions was smaller for the faster rates (task X rate interaction, F(3,84) = 5.36, p = 0.002). In the active conditions, post-hoc Bonferroni-corrected paired t-tests showed that ITPC was smaller in the 5 Hz condition relative to both the 2 Hz (t(28) = 3.51, p(corrected) = 0.009) and 3 Hz (t(28) = 4.076, p(corrected) = 0.002) conditions. No other differences between conditions were significant, although there was a trending difference where 4 Hz ITPC > 5 Hz ITPC (t(28) = 2.65, p = 0.079). In the passive conditions, no significant differences in ITPC between rate conditions were found (all p > 0.1).

The degree of separation in average neural phase between the attend high and attend low conditions (i.e. the shift in the timing of neural modulation linked to attention to one versus the other sound stream) significantly varied across rates as well (F(1,28) = 4.93, p = 0.003). Here, post-hoc Bonferroni-corrected paired t-tests showed that high-versus-low phase separation was significantly larger in the 3 Hz compared to either the 5Hz (t(28) = 3.81, p(corrected) = 0.004) or 4 Hz (t(28) = 2.90, p(corrected) = 0.043) conditions. (See **Table 1** and **Figure 2**).

**Figure 2.**
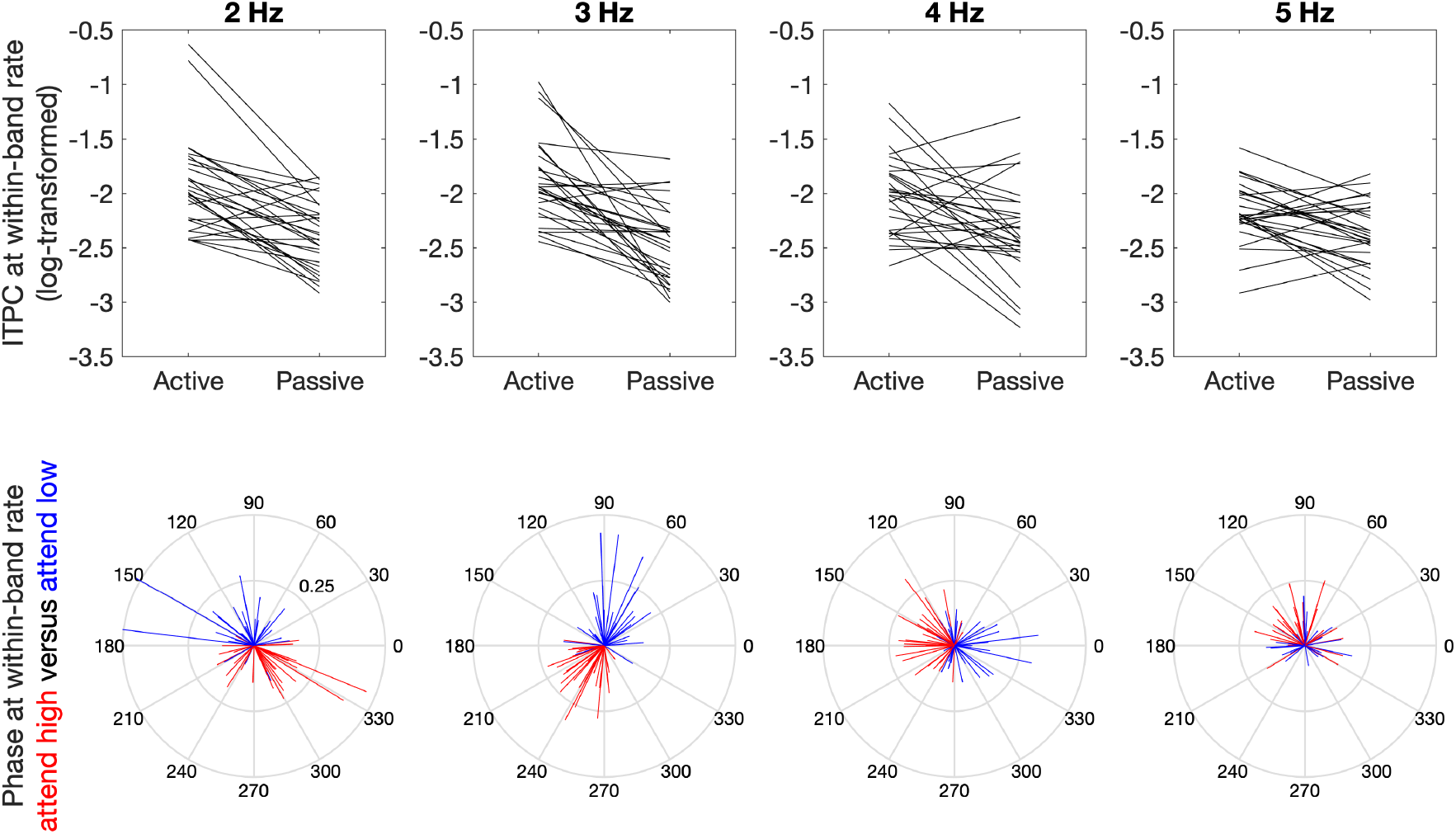
(Top) ITPC at the within-band tone presentation rate in Active and Passive conditions across four different presentation rate conditions. (Bottom) Average neural phase in the attend high (red) and attend low (blue) conditions across the four different presentation rate conditions. Each line depicts the average neural phase for a single participant; line length corresponds to the inter-trial phase coherence value for that condition; the inner grey ring marks ITPC values of 0.25, while the outer ring marks ITPC 0.50.

### EEG time-domain analyses

To investigate the effects of tone length on the passive response and attentional waveform modulation for the tone pairs, we first compared event-related waveforms for the Variable Tone Length and Fixed Tone Length Experiments, using False Discovery Rate to control for multiple comparisons (**Figure 3**). No time points survived correction for multiple comparisons, either for the comparison between passive responses, or for the attentional modulations. Importantly, there was no obvious trend for the width of the attentional modulation to be shorter in the Fixed Tone Length Experiment (in which the tones were 40 ms in length) as compared to the Variable Tone Length Experiment (in which the tone length varied between 100 and 250 ms). Given the lack of obvious differences between the responses across the two experiments, we collapsed across them in the subsequent analyses.

**Figure 3.**
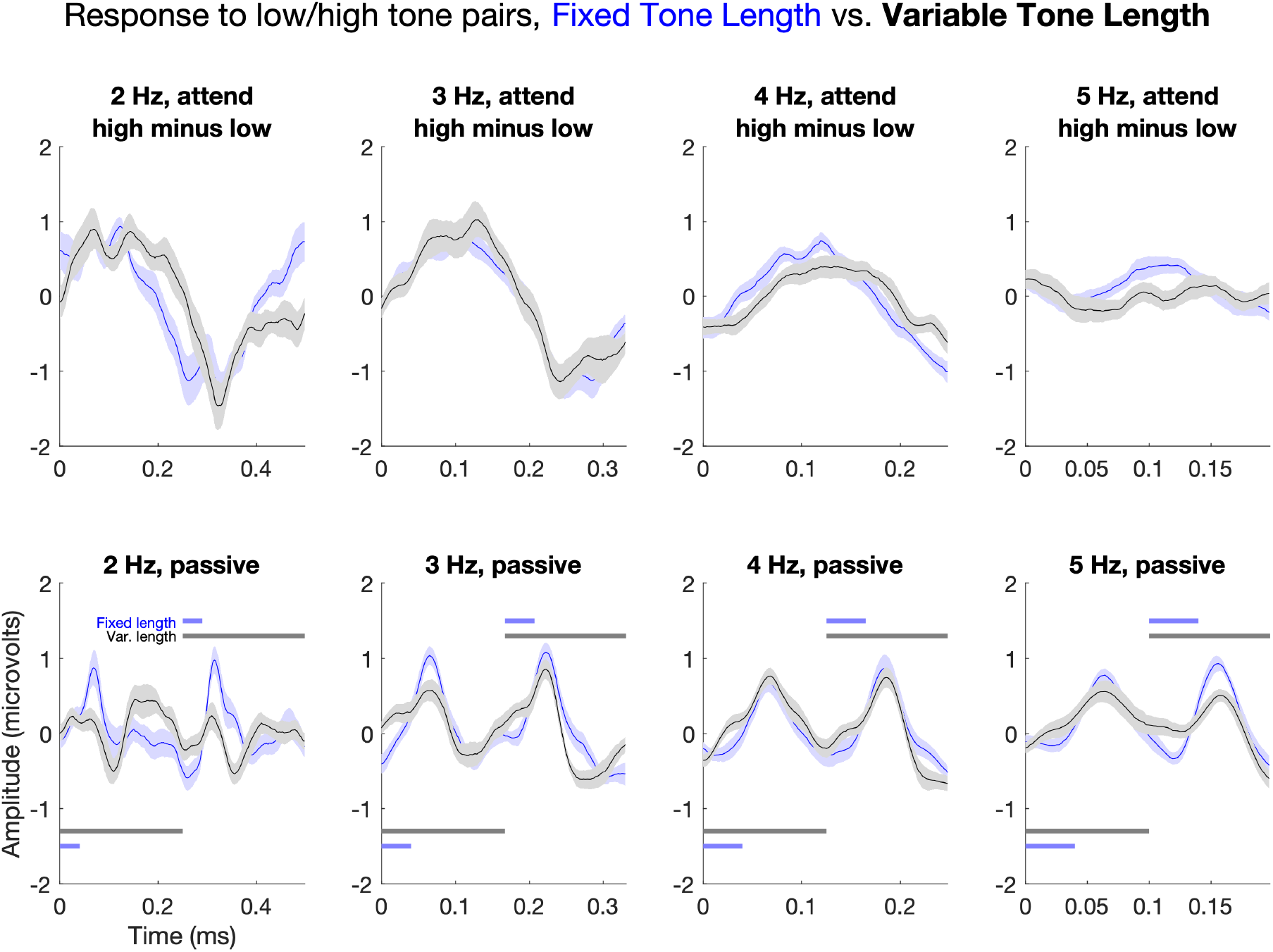
Comparison of short tone (blue) and long tone (black) waveforms across four different rate conditions. Waveforms display the average response to low-high tone pairs. Horizontal blue and gray lines display the timing of tone presentation in the fixed length and variable length conditions, respectively, with the timing of the low tone displayed at the bottom of the plot and that of the high tone displayed at the top of the plot. The top plots display the difference in waveforms between the attend high and attend low conditions, while the bottom plots display the responses in the passive conditions. Shaded regions indicate standard error of the mean. No differences between conditions survived correction for multiple comparisons.

Next, we investigated the time course of the effects by computing the difference between the responses to low-high tone pairs between attend low and attend high conditions. We first determined which portions of the response showed a significant difference across low versus high attention conditions after correction for multiple comparisons, and then analyzed the alignment between these attention effects and the shape of the passive response (**Figure 4**). Across rates, the attentional modulation followed a roughly sinusoidal shape, and the length of the modulated portions of the responses scaled with rate, such that slower rates were linked to longer modulations. For example, while in the 4 Hz condition the initial significant positive attentional modulation extended from 72 to 170 ms, in the 2 Hz condition the modulation extended from 20 to 182 ms. Across rates, the peaks of the attentional modulation aligned with the N1 of the two tones in the passive condition, such that the positive modulation peak was aligned with the N1 of the low tone response and the negative modulation peak was aligned with the N1 of the high tone response. This suggests that the attentional modulation could be partially accounted for via enhancement of the N1 response to the low tone in the attend low condition and enhancement of the N1 response to the high tone in the attend high condition.

**Figure 4.**
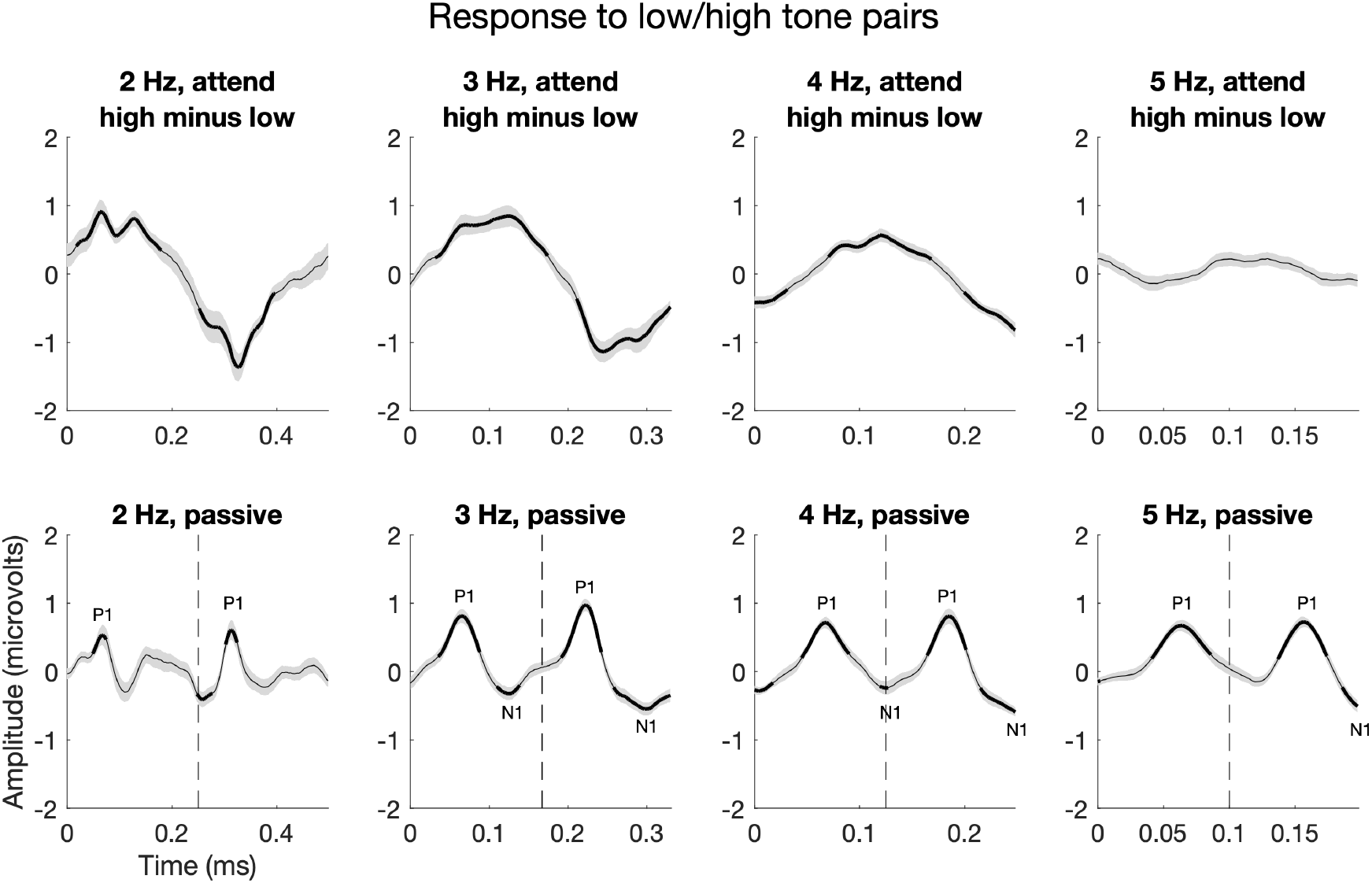
Difference between waveforms in attend high and attend low conditions (top) and passive responses (bottom) across four different rate conditions (please note difference in x-axis timescale across conditions). Waveforms display the average response to low-high tone pairs; the high tone began at the time indicated by the dotted vertical line. Shaded regions indicate standard error of the mean. Thicker lines indicate time points in which either the comparison between attend high and attend low waveforms (top) or comparison with baseline (bottom) survived correction for multiple comparisons. Note that positive-going voltage values are plotted as positive on the y-axis.

However, at the slower rates the modulation was not limited to the time points associated with the passive N1, and entirely overlapped with the time points for which the passive P1 significantly exceeded baseline. For example, at 3 Hz the initial significant positive attentional modulation extended from 33 to 176 ms, while P1 significantly exceeded baseline between 37 and 90 ms. Similarly, at 2 Hz the initial significant positive modulation extended from 20 to 182 ms, while P1 significantly exceeded baseline from 51 to 76 ms. Finally, an unexpected finding was that no significant attentional modulation was found at 5 Hz, despite the clear tone-related deflections in the passive condition.

Finally, we investigated whether effects of attention extended into the silence between sequences (beginning at the time point at which a seventh tone would have begun if the sequence were to continue through the silence). To do this, we compared the time course of these attentional modulations to the passive response, as shown in **Figure 5**. We found, in the 3, 4, and 5 Hz conditions, some evidence for a carry-over of attention effects into the between-sequence period. At 3 Hz, for example, the attend high waveform was more positive than the attend low waveform between 88 and 170 ms. This modulation closely coincided with a significant negativity in the passive waveform between 76 and 182 ms. In the 4 Hz condition, a significant negative difference was found between 0 and 55 ms, and a significant positive difference was found between 132 and 227 ms. Only the negative difference could be aligned with a significant component in the passive response (between 4 and 49 ms). Notably, at 3 and 4 Hz the time points showing significant attentional modulation overlapped with the time points showing modulations in the same direction (positive versus negative) in the response to tone pairs in **Figure 4**.

**Figure 5.**
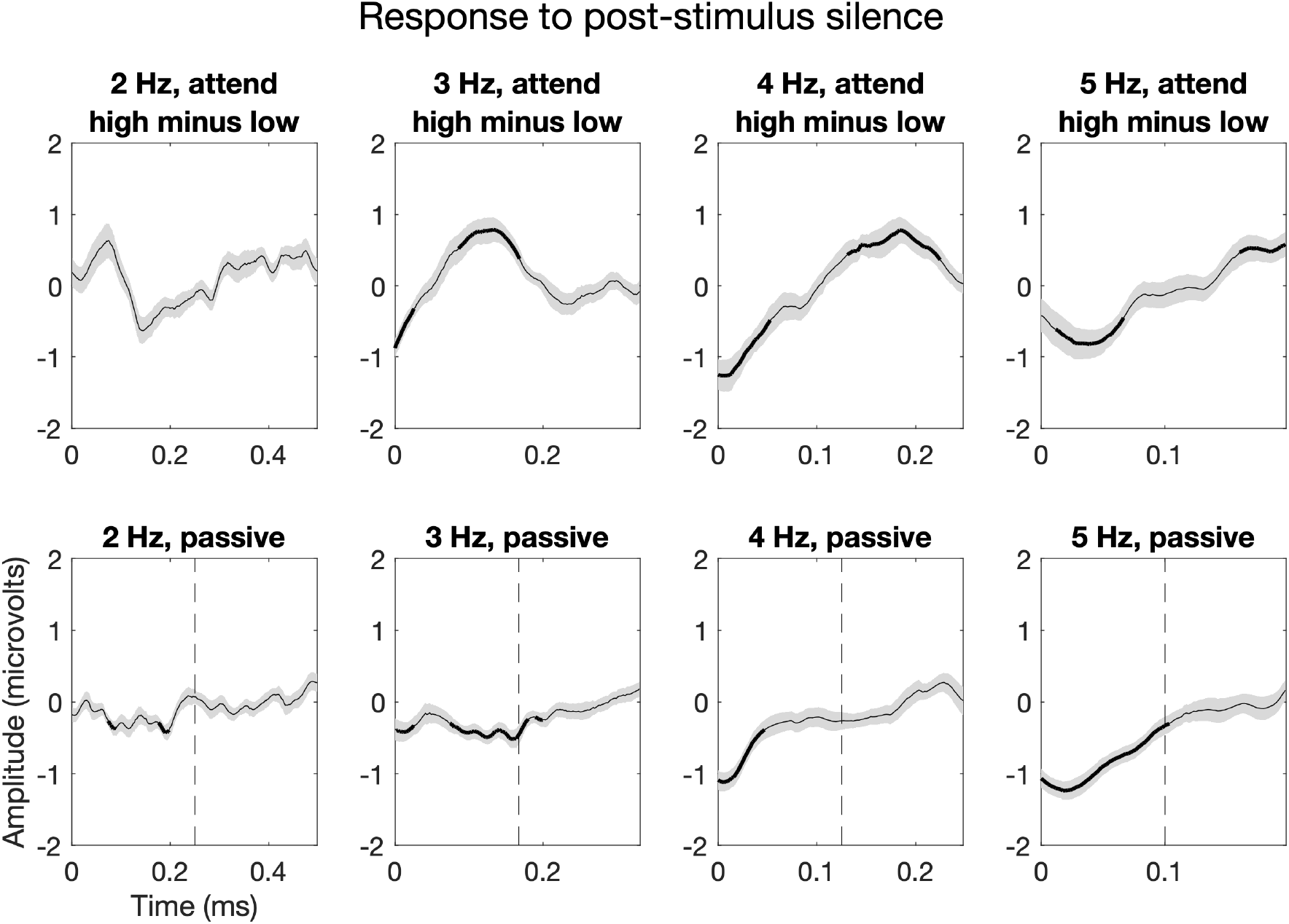
Difference between waveforms in attend high and attend low conditions (top) and passive responses (bottom) across four different rate conditions. Waveforms display the average response in the silent period between tone sequences; the dotted vertical line indicates the time at which the high tone would have sounded, had the low-high tone pairs continued through the silence. Shaded regions indicate standard error of the mean. Thicker lines indicate time points in which either the comparison between attend high and attend low waveforms (top) or comparison with baseline (bottom) survived correction for multiple comparisons.

## Summary and Discussion

Here we used a novel instantiation of an established sustained auditory selective attention task - where participants attend to one of two sound streams presented concurrently at the same rate but at opposite phases - to adjudicate between two general theoretical accounts of auditory attentional mechanisms: an endogenous neural entrainment account versus an attentional gain account. Specifically, we asked whether attention-driven phase-locking reflects alignment of slow endogenous neural rhythms with the temporal structure of the stimuli, versus attentional modulation (or ‘gain’) of evoked waveforms. To disambiguate between these theoretical accounts, we manipulated both tone stream rate (within experiment) and tone duration (across experiments). Under the endogenous attentional entrainment account, we would expect that a) the width of the phasic attentional modulation would scale inversely with increasing rate; b) phasic attentional modulation width would remain relatively insensitive to changes in tone duration; and c) phasic attentional modulation would continue into the silent period at the end of each trial. By contrast, on an attentional gain account, we would expect that a) the width of attentional modulation would remain relatively stable across tone presentation rates; b) attentional modulational width would increase with longer tone duration; and c) the phasic attentional EEG waveform would not continue into the silent period of each trial.

We found, when comparing the evoked waveforms elicited when attention was directed towards one versus the other stimulus stream, that direction of attention was linked to a roughly sinusoidal pattern of periodic positive and negative modulations. These negative/positive modulations were temporally centred on the N100 evoked responses to each of the tones in the attended/ignored stream, as recorded in the passive condition. This finding is broadly consistent with prior reports that the N100 can be modulated by selective attention (Hillyard et al., 1973; Sanders & Astheimer, 2008; Choi et al., 2013; Dai et al., 2018). However, the width of the attentional modulations scaled with rate, such that slower rates were linked to wider attentional modulations. Thus, at slower rates the modulation extended into the time regions associated with other responses in the passive condition, including the P50. The only way this pattern of attentional modulations could be accounted for via modification of evoked exogenous responses would be to suppose that the P50 was suppressed by attention. However, prior research suggests that the P50 is, rather, enhanced by attention (Woldorff & Hillyard, 1991; Woldorff et al., 1993).

Our finding that, at slower rates, the increased negativity associated with selective attention extends outside of the time region containing the N100 and into the time region containing the P50 suggests that modulation of exogenous responses cannot be the primary mechanism underlying the attention-driven changes in phase locking. Moreover, our finding that the width of the modulation of the evoked waveform by attention was similar in the Fixed Tone Length and Variable Tone Length experiments suggests that our results cannot primarily reflect modulation of an amplitude-envelope-following response. However, this does not mean that the involvement of neural oscillations must necessarily be invoked to explain our findings. An alternate possibility is that the results reflect modulation of an endogenously generated potential that, at slower rates, exceeds the width of the N100. For example, a number of studies investigating the effects of selective attention on ERPs have reported the existence of a prolonged negative shift (sometimes labelled Nd) that is temporally dissociable from the N100 (Näätänen et al., 1978; Michie et al. 1990; Woods & Alain, 1993, 2001; Degerman et al., 2008). Thus, our results could arguably be accounted for by invoking modulation of Nd. However, the Nd has not previously been reported to scale with rate in a manner that could generate the pattern of modulations across conditions which we find. For example, Woods and Alain (1993) report an Nd lasting around 200 msec in an experiment using inter-onset-intervals of only 50-210 milliseconds, whereas we found that in the 4 Hz condition (corresponding to 250 millisecond inter-onset intervals) the duration of the modulation was less than 100 ms. Moreover, even if our results could be explained via the modulation of endogenous evoked potentials with a duration that scales with rate in a linear manner, generating roughly sinusoidal difference waves between attention conditions, it is not apparent whether this explanation differs substantially from the hypothesis that sinusoidal low-frequency neural activity phase-locks to the temporal structure of attended stimuli.

If attentional modulation of neural entrainment reflects the alignment of endogenous neural oscillators with the temporal structure of attended stimuli, then attention-driven neural modulations should continue for a time even once stimuli have ceased, given that oscillations are theoretically somewhat self-reinforcing (Large & Kolen 1994). To test this hypothesis, we designed the task such that participants attended to a series of short tone sequences so that we could investigate whether the sinusoidal attentional modulation continued through the silence between sequences. The evidence regarding continuance of oscillations was somewhat equivocal. In the 3 Hz and 4 Hz conditions, modulations were present in the silence between conditions, and there was temporal overlap between the modulated portions of the silence and the modulated portions of the phase cycle during stimulus presentation. Moreover, in the 5 Hz condition we found a significant modulation present during the silence despite finding no significant modulation during stimulus presentation. However, for the most part these modulations overlapped closely with a late negative potential that was also present in the passive condition, suggesting that they could simply reflect modulation of the N2. Furthermore, we found no significant continuation of attentional modulations into the silence in the 2 Hz condition, even though in this condition modulations were robust during the sequence.

An unexpected finding was that there were no significant modulations of the evoked waveform in the 5 Hz condition, suggesting that attentional modulation of neural entrainment differed across rates. To investigate this possibility, we compared effects of attention on ITPC and neural phase across rate conditions. The difference in ITPC between attention and passive conditions was significantly smaller in the 5 Hz condition compared to the 2 Hz and 3 Hz conditions, while the difference in phase between the attend high and attend low tasks was significantly smaller in the 5 Hz condition compared to the 3 Hz and 4 Hz conditions. The lack of attentional modulations in the 5 Hz condition could partially reflect task performance, which was lower in this condition compared to the other three rates. However, performance was still well above chance in the 5 Hz condition, and this paradigm has been shown to be sensitive to the neural effects of selective attention even in participant populations who display relatively poor performance (such as children with ADHD; Laffere et al., 2020b). The enhanced attentional modulations at slower rates could partially reflect the well-known dominance of low frequencies in the EEG signal (Näpflin et al., 2007); however, there was no difference in ITPC between the 4 rate conditions for the passive task.

The relative lack of attentional modulation in the 5 Hz condition compared to the other rates could reflect a switch from a temporally-selective task strategy to a spectrally-selective strategy. Prior research has suggested involvement of the motor system in perception of an isochronous beat underlying a complex rhythmic stimulus, with activation of cortical and subcortical areas found during beat perception tasks even when participants were explicitly instructed not to move (Grahn & Brett, 2007; Grahn & Schuit, 2012). One possibility, therefore, is that the motor system generates temporal predictions about the onset of an attended sound stream (Morillon & Schroeder, 2015). Temporal predictions which rely on the motor system may be rate-limited, as individuals’ ability to align movements with the temporal structure of sound streams falls off rapidly as the stimulus presentation rate is increased (Repp, 2003). Supporting the involvement of implicit motor movement in temporal attention, Zalta et al. (2020) found that both motor tapping and auditory temporal attention showed similar dependence on rate, with optimal performance at close to 2 Hz and poorer performance at faster and slower rates. The involvement of the motor system in temporally-selective attention is also supported by our previous finding that performance on the non-verbal selective attention task and the effect of attention on neural phase-locking correlated with performance on tests of synchronization and rhythm reproduction (Laffere et al., 2020b).

In conclusion, we find that attention to one of two sound streams, presented at the same rate but out of phase, is linked to an increase in phase-locking at the presentation rate and a shift in neural phase between attention conditions. At slower rates, the attentional modulation of the evoked waveforms dissociates in time from passive evoked potentials, suggesting that the changes in phase alignment reflect synchronization of slow endogenous neural activity with the temporal structure of the attended stimulus rather than attenuation or enhancement of exogenous responses to stimuli. However, these modulations became smaller as the presentation rate increased, vanishing entirely once the rate reached 5 Hz. This suggests that neural entrainment may only be a useful strategy for attentional selection at slower presentation rates, and that listeners may rely upon alternate mechanisms at higher rates.

## Supporting information

Supplemental example stimuli

## Notes

***Conflicts of interest*:** The authors have no competing interests to declare, financial or otherwise.

### Competing Interest Statement

The authors have declared no competing interest.

